# A global Ramachandran score identifies protein structures with unlikely stereochemistry

**DOI:** 10.1101/2020.03.26.010587

**Authors:** Oleg V. Sobolev, Pavel V. Afonine, Nigel W. Moriarty, Maarten L. Hekkelman, Robbie P. Joosten, Anastassis Perrakis, Paul D. Adams

## Abstract

Ramachandran plots report the distribution of the (φ, Ψ) torsion angles of the protein backbone and are one of the best quality metrics of experimental structure models. Typically, validation software reports the number of residues belonging to “outlier”, “allowed” and “favored” regions. While “zero unexplained outliers” can be considered the current “gold standard”, this can be misleading if deviations from expected distributions, even within the favored region, are not considered. We therefore revisited the Ramachandran Z-score (Rama-Z), a quality metric introduced more than two decades ago, but underutilized. We describe a re-implementation of the Rama-Z score in the Computational Crystallography Toolbox along with a new algorithm to estimate its uncertainty for individual models; final implementations are available both in Phenix and in PDB-REDO. We discuss the interpretation of the Rama-Z score and advocate including it in the validation reports provided by the Protein Data Bank. We also advocate reporting it alongside the outlier/allowed/favored counts in structural publications.

## Introduction

Validation is an integral part in obtaining three-dimensional models of macromolecules in X-ray crystallography (MX; Read *et al*., 2011) and in cryo-Electron Microscopy (cryo-EM; Henderson *et al*., 2012). It is also key in interpreting the quality of models from the Protein Data Bank (PDB; Burley *et al*., 2019), as there is no formal structure quality requirement for acceptance to this repository. A key quality metric used in validation of the quality of atomic models of proteins is the Ramachandran plot (Ramachandran *et al*., 1963). Ramachandran plots describe the two-dimensional distribution of the protein backbone (φ, Ψ) torsion angles. They have been used for the validation of protein backbone conformations since their introduction in PROCHECK (Laskowski *et al*., 1993), and then later in software packages such as O (Kleywegt & Jones, 1996), WHAT_CHECK (Hooft *et al*., 1996) and MolProbity (Lovell *et al*., 2003). The phrase “no Ramachandran plot outliers” is widely considered as the “gold standard” for a high-quality structure and is often found in the main text of papers reporting protein structures, while the absolute number or the percentage of residues in the so-called “outlier”, “allowed” and “favored” regions is typically reported in tabular form. It should be noted that a better phrase is “no *unexplained* Ramachandran plot outliers”, as it is not uncommon for there to be a very small number of legitimate outliers in the plot, which are supported by the experimental data and often relate to some functional aspect of the protein (Richardson *et al*., 2018).

All software for refining macromolecular models uses a standard set of stereochemical restraints on covalent geometry (Engh & Huber, 2012): these provide sufficient information for structures at 3.0Å resolution or better. Advances in electron cryo-microscopy (Li *et al*., 2013; Bai *et al*., 2015) have led to greatly improved resolution of cryo-EM maps, but while this improved resolution enabled full-atom refinement of macromolecular structures (Afonine *et al*., 2018; Nicholls *et al*., 2018), the majority of cryo-EM models are still solved in the 3-5Å resolution range. Likewise, in X-ray crystallography, low-resolution data sets remain an issue: atomic modeling and refinement against low-resolution data is challenging and can benefit substantially from using all available a priori knowledge about the molecule at hand (Kleywegt & Jones, 1998).

At low resolution it is often necessary to use information beyond the stereochemical restraints on covalent geometry: internal molecular symmetry (Kleywegt, 1996); homologous structure models determined in higher resolution as a reference (Smart *et al*., 2012; Nicholls *et al*., 2012; Headd *et al*., 2012; Schröder *et al*., 2010) or as a source for hydrogen bond length restraints (Beusekom *et al*., 2018); information about secondary structure and rotameric states of protein amino-acid side chains (Headd *et al*., 2012) have all been used in various software implementations to provide additional information in low resolution refinement. Clearly, the well-defined distribution of protein main-chain φ and Ψ angles in Ramachandran space is yet another source of information that can guide model building and refinement (Kleywegt & Jones, 1996). Ramachandran restraints can help prevent deterioration of backbone conformation during low-resolution refinement, thereby maintaining chemically meaningful model stereochemistry. Many software packages provide an option to use Ramachandran restraints, e.g. in XPLOR/CNS (Brunger, 1993; Brünger *et al*., 1998), QUANTA (Oldfield, 2001), Coot (Emsley *et al*., 2010) and Phenix (Headd *et al*., 2012). The Rosetta (Leaver-Fay *et al*., 2011) all-atom energy function includes term based on Ramachandran distribution (Alford *et al*., 2017).

While helpful for refinement, actively using the Ramachandran plot as a source of restraints reduces its utility as an independent validation metric. It can, however, still be used to report on the model quality, similar to how bond length and bond angle deviations are reported even though these are nearly always restrained. A larger issue arises during the construction of Oldfield-like (Oldfield, 2001) Ramachandran restraints: assigning (and restraining) each pair of (φ, Ψ) angles in the model to a nearby target within the plot. This assignment is purely reliant on the input model, which may not be correct. Incorrectly fit peptide planes, which are associated with large differences between the current and the correct position of two residues on the Ramachandran plot, occur in more than half of all atomic models (Joosten *et al*., 2012) and we have previously shown that correction of such errors requires model rebuilding (e.g. peptide flipping) rather than refinement (Joosten *et al*., 2011). Errors in starting models will lead to incorrect Oldfield-like restraint target assignments and (φ, Ψ) errors will propagate into the model as a result of refinement (Kleywegt & Jones, 1998; Richardson *et al*., 2018). The refined model may then appear to have desirable Ramachandran statistics in terms of expected fractions of residues belonging to favored/allowed/outlier regions, while the distribution of (φ, Ψ) itself is improbable. Unfortunately, this may not be obvious to an untrained eye.

We illustrate this in Figure 1, contrasting a nearly perfect-looking Ramachandran plot^1^ derived from an ultra-high resolution structure (Fig. 1, left) with an obviously poor plot (Fig. 1, middle). The ultra-high resolution plot in Figure 1, left contains few outliers, has a majority of points in the favored region and follows the observed distribution that is shown in the background color of the plot. The middle plot in Figure 1 has a preponderance of outliers, and is trivial to identify both visually and by the number of outliers. In Figure 1, right, however, we illustrate that simple visual inspection or outlier count can be misleading: while residues are within the most favored region with no outliers, the distribution does not coincide with the most favorable peak (darkest blue) in alpha-helical and beta-sheet regions. While such anomalies are apparent to a trained eye, they are not reflected either by the count or the percentage of outliers, nor in the counts in outlier/allowed/favorable regions: these numbers are practically identical between figures 1 (left) and 1 (right) making it hard both to computationally and manually mine for such anomalies to report them in a clear manner.

**Figure 1.**
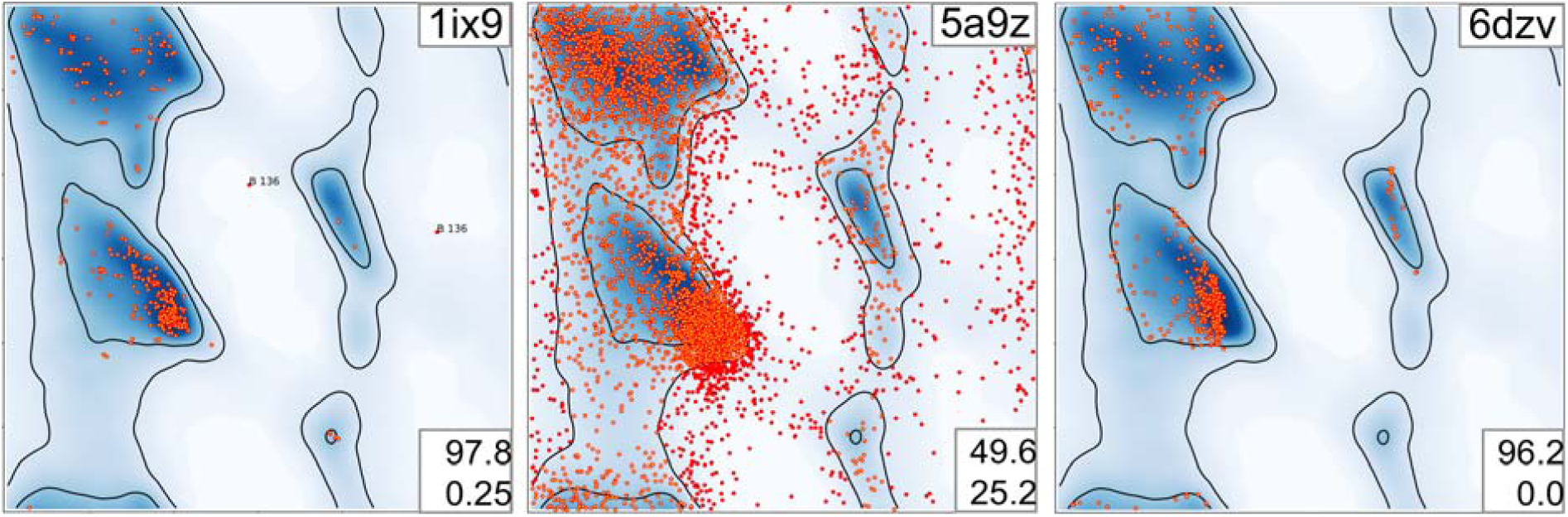
Examples of Ramachandran plots: Left: a good-looking Ramachandran plot for (1ix9, 0.9 Å), Middle: an obviously bad Ramachandran plot (5a9z, 4.7 Å) and Right: a suspicious Ramachandran plot (6dzv, 4.2 Å). PDB ID code of the models in top right corner. Two numbers (bottom right) indicate percentage of residues in favored (top) and outlie (bottom) regions.

Fortuitously, (Hooft *et al*., 1997) proposed a numerical metric, called the Ramachandran Z-score (Rama-Z), that characterizes the shape of the (φ, Ψ) angle distribution in the Ramachandran plot. That metric was based on the statistical analysis of high-quality models available in the PDB at that time. While this metric has been available since 1997 in the WHAT_CHECK program and has been routinely reported by PDB-REDO (Joosten *et al*., 2009, 2014), it never made it into mainstream validation pipelines (Read *et al*., 2011) nor did it become standard practice to report the metric in the model quality summary in “Table 1”. There is now an avalanche of lower resolution structures being deposited, in a large part thanks to the cryo-EM revolution (Li *et al*., 2013; Bai *et al*., 2015), and to dramatic improvements in refinement methods (Afonine *et al*., 2018; Nicholls *et al*., 2018) that now actively exploit the majority of available a priori information about model geometry. The explicit use of the (φ, Ψ) angles distribution in refinement now makes it essential for new model quality measures that are independent of the information used in the refinement target function.

Here we demonstrate the value of this old but powerful Rama-Z validation metric, and showcase its utility across a number of examples where standard validation tools fail to pinpoint the issue. We describe the implementation of the Rama-Z score in CCTBX (Grosse-Kunstleve *et al*., 2002) and propose a method to estimate its reliability for a particular model. Implementations taking into account the current distribution of (φ, Ψ) angles in high-quality and high-resolution models in the PDB and including the reliability metric are now available both in Phenix and PDB-REDO. Based on specific examples and a global evaluation we argue that the Rama-Z score should be broadly adopted.

## Results and discussion

### Interpretation of the Rama-Z score

The Rama-Z score describes how ‘normal’ a model is compared to a reference set of high-resolution structures (Hooft *et al*., 1997). As in the original paper, we calibrated the score to make the average score for the reference set zero with a standard deviation of 1.0. The original paper suggested that a Z-score of −4 and lower indicates a serious problem with the structure. We suggest stricter cutoffs since the number of models in the reference set is significantly bigger and we can expect that the distributions will account for rarer but still valid cases. Large positive Rama-Z scores also show significant deviation from the reference distribution hence they are as unlikely as large negative ones. Presuming a normal distribution, only 0.2% of structures would be expected to have |Rama-Z| > 3; however, we observe that only 0.04% of high-resolution models have |Rama-Z| > 3 in the PDB. Therefore Rama-Z scores with absolute values above 3 correspond to geometrically improbable structures (in terms of main chain geometry), absolute values between 3 and 2 (which would encompass 4.2% in a normal distribution and in practice are 0.8% of high-resolution models) correspond to less likely yet possible models, and anything between −2 and 2 indicates normal protein backbone geometry.

The Rama-Z score is a global metric that provides an overall assessment of model quality and is not able to report on local issues with the main chain conformation. We also note that apart from the single, global, Rama-Z score, separate Rama-Z scores are calculated for strands, helices, and loops: these are worth checking, especially for “suspicious” Rama-Z values (2<|Rama-Z|<3). The single value of Rama-Z on its own is still most useful for an overall assessment of the model. However, we recognize that some applications would require an estimate of the reliability of the Rama-Z score for the model being analyzed. An application that requires tracking both the Rama-Z score and its RMSD is the rapid evaluation of whether the backbone geometry is significantly better after rebuilding and refinement, as performed in PDB-REDO. When the overall model quality and fit to the experimental data improves, the Rama-Z score typically follows, and can be used to show model improvement in PDB-REDO runs (Joosten *et al*., 2009). The RMSD can also be used to assess the significance of a difference in Rama-Z score between two models; an implementation of this feature in the context of PDB-REDO is shown in Suppl. Figure 2. As the reliability of the Rama-Z score was not explored in the original paper, we developed a method for its calculation and call it Rama-Z RMSD (RMSD for short from herein; see Methods).

**Figure 2.**
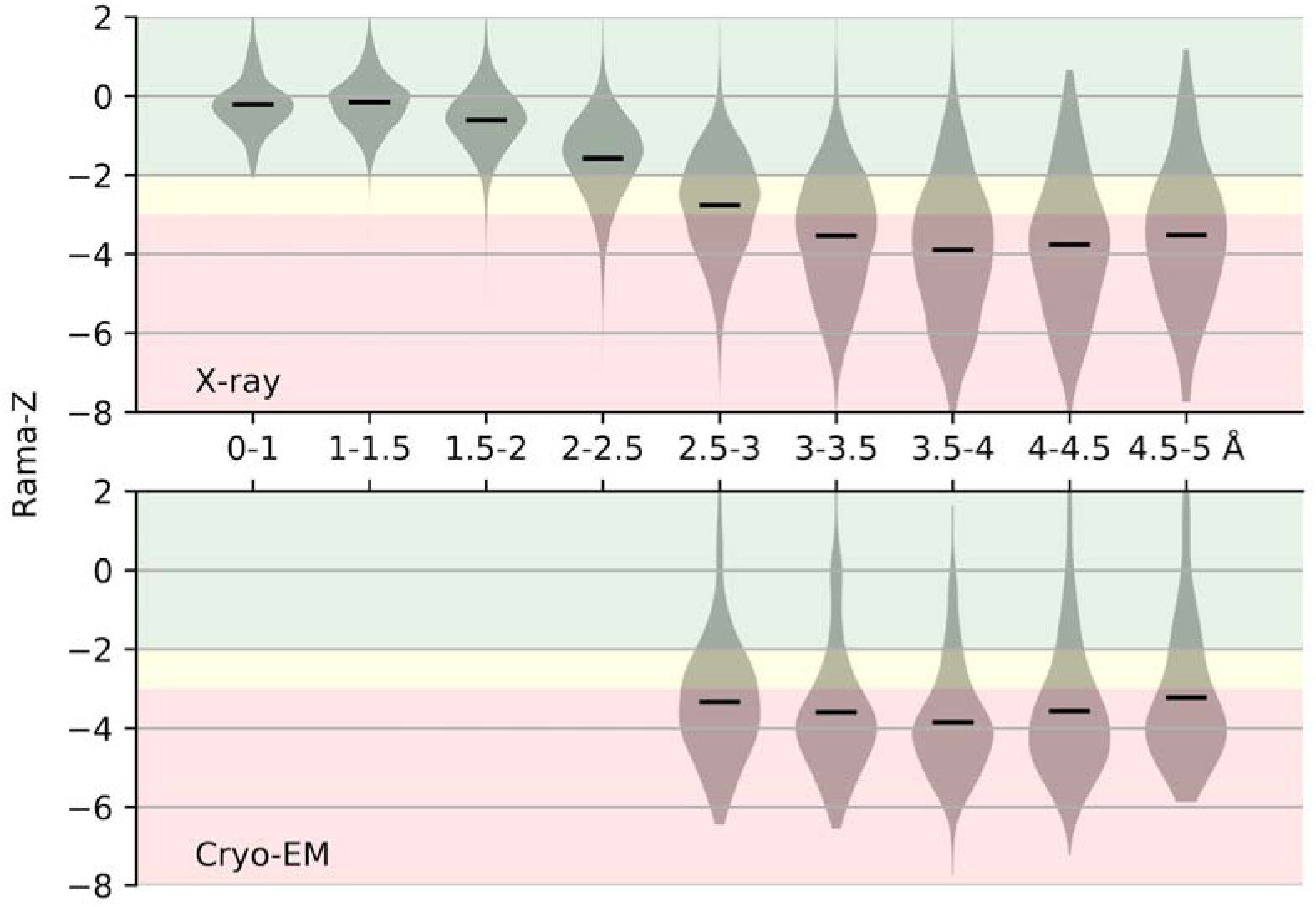
Distribution of Rama-Z scores by resolution for structures solved by X-ray diffraction and cryo-EM. Solid horizontal bars on each violin indicate the mean. The background color represents proposed Rama-Z ranges. Red (below −3 and above 3) is for geometrically improbable backbone geometry, yellow (from −3 to −2 and from 2 to 3) for unlikely yet possible, green (from −2 to 2) is for normal backbone geometry.

### Rama-Z scores for models in the Protein Data Bank

We calculated the Rama-Z score for all X-ray (107,800) and cryo-EM (1,711) structures available in the PDB, (Figure 2). The vast majority of the models at resolution of 2Å or better have Rama-Z scores that we consider good (between −2.0 and 2.0); most models between 2-2.5Å resolution also have good Rama-Z scores. While there is a clear trend that the Rama-Z score distribution deteriorates at lower resolution, rather surprisingly, the distribution and median between 3 and 5Å remain almost constant. Slightly better values for very low-resolution models are likely a result of these models being complex structures of fitted higher-resolution homologous models and do not correspond to de-novo modeling and atomic refinement against the data. The distribution for cryo-EM models is similar to X-ray models at matching resolutions, with the only observable trend being better models for X-ray crystallography between 2.5 and 3.0Å.

In Figure 3 we show the Rama-Z score and the percentage of residues in the favored regions of the Ramachandran plot for cryo-EM and X-ray structures solved at resolutions better than 5Å. We filtered out small structures with fewer than 100 protein residues because their Rama-Z values usually have a large uncertainty (see Methods), leaving 104,470 structures. It is clear that the Rama-Z score correlates with the fraction of residues in favored regions of the Ramachandran plot. Among models solved with experimental data in the 1.2-5Å range (black dots in Figure 3) 28% have Rama-Z < −2 and 14% have Rama-Z < −3; 0.19% have Rama-Z > 2, and only 0.01% have Rama-Z > 3. At the same time, of the high-resolution models (better than 1.2Å, dark blue dots on the plot) only 0.4% have Rama-Z < - 2 and no model has Rama-Z < −3. Similarly, 0.4% have Rama-Z > 2 and 0.04% have Rama-Z > 3. The red “x” in Figure 3 denotes structures with a relatively high percentage of Ramachandran favored residues, but low Rama-Z score. These examples are discussed in more detail below (Selected examples from cryo-EM) and the corresponding Ramachandran plots are shown in Figure 4.

**Figure 3.**
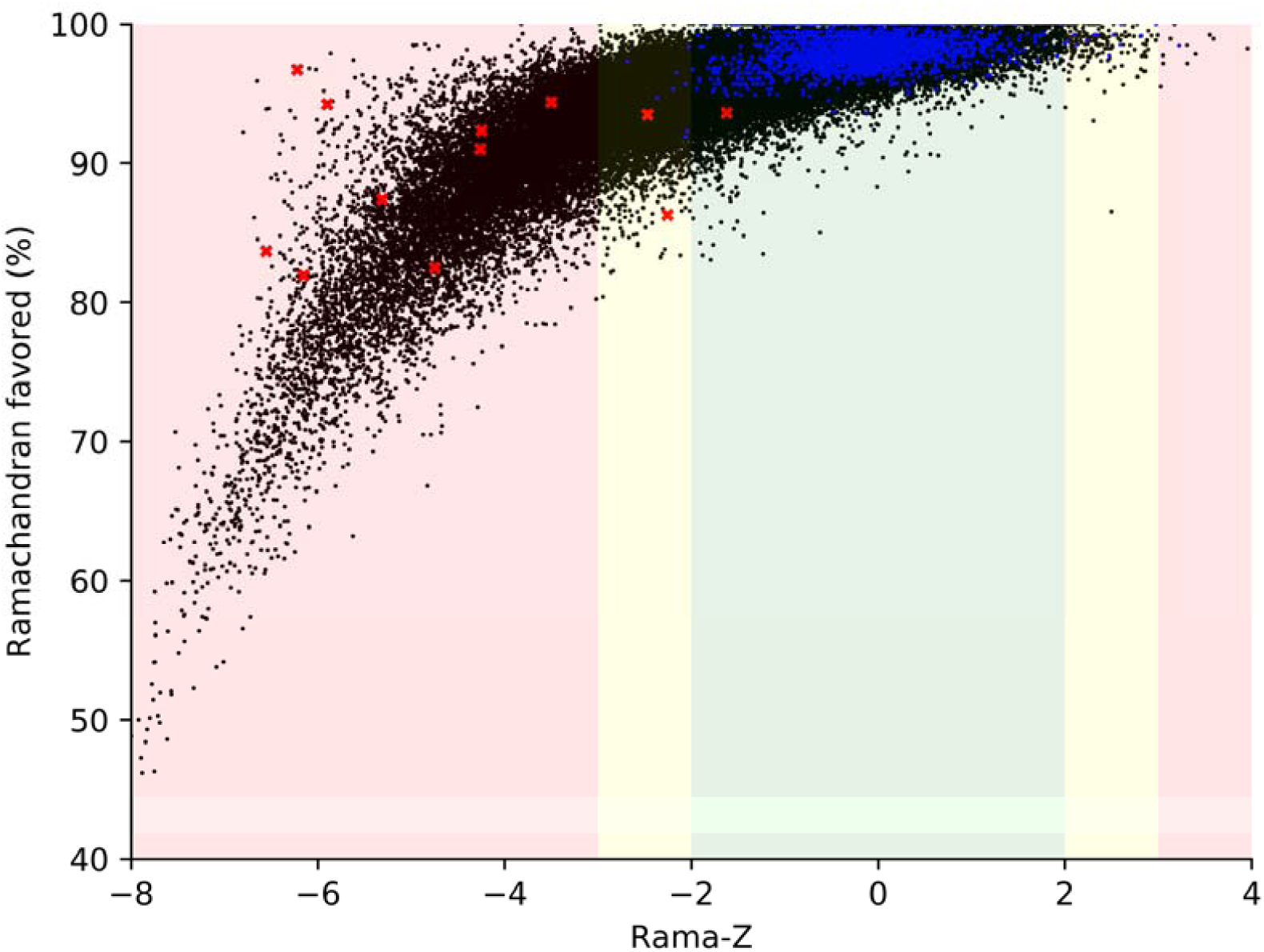
Rama-Z values versus percentage of residues in favored region of Ramachandran plot for all X-ray and cryo-EM derived models in PDB at resolution 5Å or better containing more than 100 amino acid residues. Blue dots represent models with resolution better than 1.2 Å, black dots represent models with resolution between 1.2-5 Å, red crosses represent models shown on Figure 4. The background color represents proposed Rama-Z ranges. Red (below −3 and above 3) is for geometrically improbable backbone geometry, yellow (from - 3 to −2 and from 2 to 3) for unlikely yet possible, green (from −2 to 2) is for normal backbone geometry.

**Figure 4.**
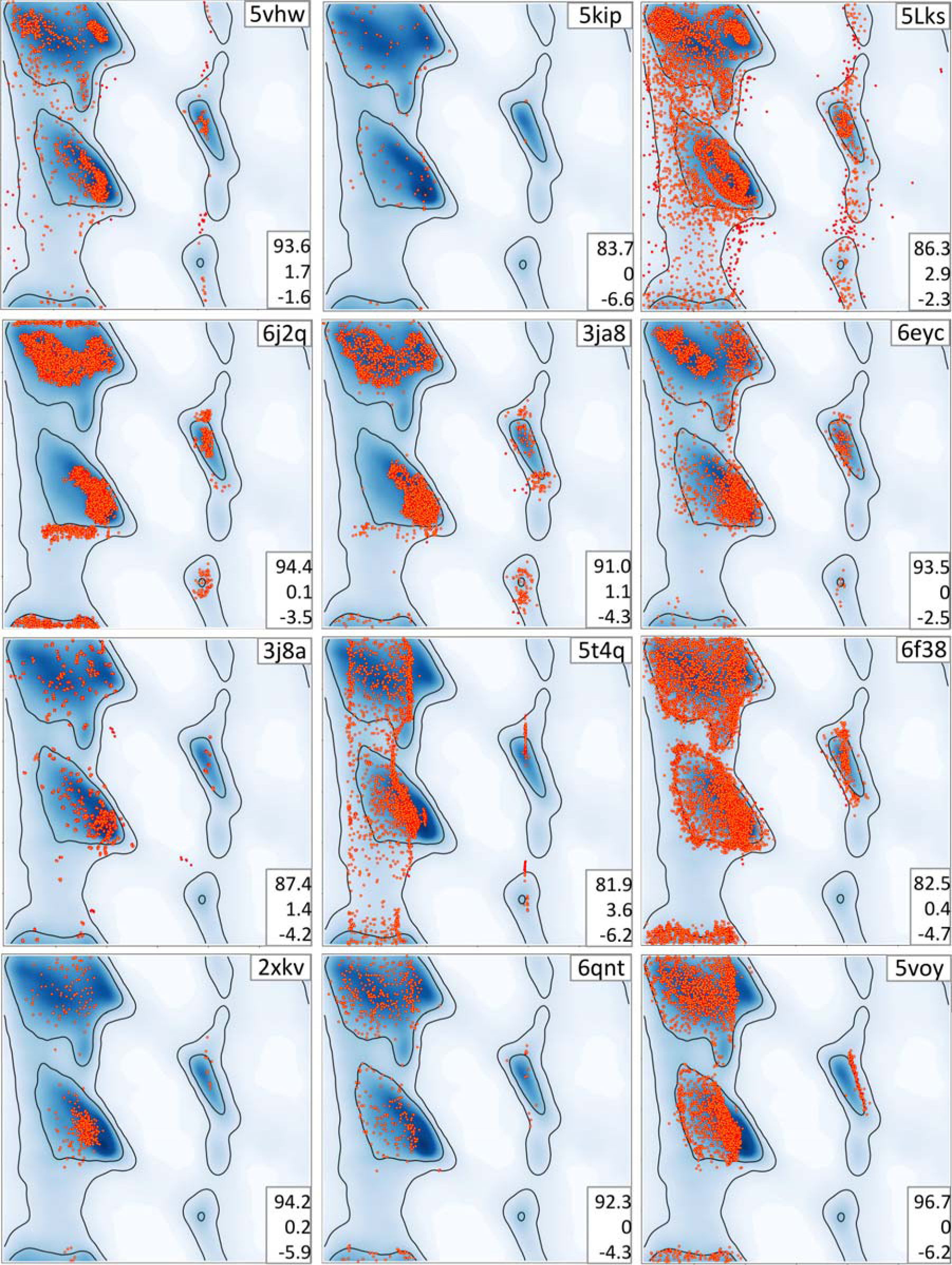
Examples of Ramachandran plots with unusual distribution of (φ, Ψ) angles. Plots are referred to by the PDB code of corresponding atomic model (upper right corner of each plot). Triplets of numbers on the bottom right on each plot indicate, from top to bottom: percentage of residues in favored and outlier regions, Rama-Z.

### Selected examples from cryo-EM

We here analyze in more detail twelve examples of problematic plots, where the underlying issues are less obvious (red crosses in Figure 3 and more detail in Figure 4). All the models in these examples are from cryo-EM; with most in the 3.5-4.0Å resolution range. They are not single occurrence examples, but are rather representatives of whole sets of models with similar artifacts of the Ramachandran plot. Below we discuss the pathologies in these structures that are trivial to detect using the Rama-Z score (all but three examples have Rama-Z < −3.0) but could go unnoticed using the standard favored or outlier metrics (many examples have >90% in the favored region and all but one example have less than 2% outliers).

#### 5vhw

In this 7.8Å structure, except for a few outliers, the plot itself does not appear very unusual, except maybe for slightly systematic clustering of residues in the alpha and beta regions, although this does not produce a particularly alarming Rama-Z score (−1.6). However, the individual Rama-Z values are −1.9, −2.5 and 0.7 for helices, sheets and loops respectively. The low score for the beta-sheet part of the model explains the somewhat odd-looking distribution of residues in the beta region. Although the helix score of −1.9 belongs in the good range, it borders on “suspicious”, which is again in line with the somewhat unusual distribution in the alpha region.

#### 5kip

All residues belong to the favored and allowed regions (no outliers) in this 3.7Å structure. Visually the plot does not trigger any major alarm, except that the residues look almost randomly distributed inside the favored regions with only 83.7% inside these regions. A very low Rama-Z score highlights this oddity.

#### 5Lks^2^

This 3.6Å structure shows one of the most usual looking plots in this series, with various strangely shaped ridges separated by nearly empty valleys in the favored regions. A Rama-Z score of −2.3, in the suspicious range, raises concerns. Individual scores of −4.0, −4.2 and 0.2 for helices, sheets and loops provide the insight that poor distributions and scores for helices and sheets are masked by the overall good distribution for the loops. Notably, 65% of this this model consists of loops but this was not enough to completely suppress the overall Rama-Z score for reasons described in Methods.

#### 6j2q

The plot for this 3.8Å structure shows residues clustering exactly around peaks of the favored regions and two peculiar horizontal clusters: one in the allowed region just below the helices and one at the favored strands region (including residues both on the top and on the bottom of the plot); the Rama-Z score clearly identifies this issue, with a low overall score of −3.5.

#### 3ja8 and 6eyc

these two entries represent the same model at 3.8Å resolution; 6eyc a version of 3ja8 that has been extensively re-built and re-refined manually and made available as a separate entry in the PDB (Croll, 2018). The plot for the rebuilt structure (6eyc) is much improved, as clearly indicated by the Rama-Z score (−2.5 instead of −4.3). Although 6eyc still shows an unusual distribution in the helices and particularly in the strand region (note the three distinct clusters to the left), the Rama-Z score does not report it as poor but rather *suspicious*.

#### 3j8a

In this 3.8Å structure, the number of residues in Ramachandran favored region is already low (87.4%, lower than the MolProbity guidelines of 90%) and that is also evident in the Rama-Z score (−4.2). That alone hints at some problems with backbone geometry. Additionally, these residues are clustered in groups of four residues due to the presence of chains related by internal molecular symmetry.

#### 5t4q

This very low-resolution structure (8.5Å) provides another unusual looking plot. The residues form essentially two clusters: one broad lane on the left, spanning the alpha and beta regions and one very slim line on the right. This is very unlikely to represent the real main-chain conformation of the protein and this is clearly highlighted by the very low Rama-Z score of −6.2. Such a distribution is very likely created by the use of an inappropriate target function in the model’s refinement and the low resolution of the map itself which doesn’t justify refinement of atomic coordinates.

#### 6f38

In yet another low-resolution structure (6.7Å) this is another instance of a very unlikely distribution. The residues lie along the borders of the alpha and the beta regions; no residues are observed in between the regions. The Rama-Z score highlights this with a poor score of −4.7. This is likely a result of refinement (or pure geometry optimization) using strong Ramachandran plot restraints, starting with a model that had many Ramachandran plot outliers.

#### 2xkv, 6qnt and 5voy

These three models, at resolutions of 13.5, 3.5 and 7.9Å respectively, have the majority of residues in favored regions, with virtually no outliers. However, we note that all residues are distributed such as to completely avoid the most prominent peaks of the plot. This typically happens when stereochemistry terms with a strong non-bonded repulsion dominate refinement target and the model does not include explicit hydrogen atoms. All three cases receive very poor Rama-Z scores.

### High values of Rama-Z score from cryo-EM

Very low values of Rama-Z score indicate unlikely backbone geometry and probable artifacts in the Ramachandran plot. At the same time, very high values may also indicate unusual Ramachandran distributions and sub-optimal backbone geometry. One example is illustrated in Figure 5. Two 100% sequence identical structures are considered: 1jz7 was solved by X-ray crystallography at 1.5Å resolution and has a good Rama-Z score of −0.8, while 3j7h was solved by cryo-EM at 3.2Å resolution and has Rama-Z score of 2.4. Both models are very similar, with a root-mean-square deviation between the main-chain atoms of 0.65Å. However, as the 1jz7 model has an excellent fit (Rfree 0.22) to high-resolution data, it is much more likely to represent the true structure. The high Rama-Z value of the lower resolution cryo-EM model (3j7h) indicates problems, and this is reinforced by a MolProbity clashscore of 131. Indeed, the researchers used tight Ramachandran restraints in Coot for the entire model optimization process (Bartesaghi *et al*., 2014). This resulted in residues clustering almost exclusively on top of Ramachandran plot peaks, which is a very unlikely distribution. This serves as a warning that one should be careful to avoid over-optimizing, especially using unreasonable weights in restraints such as those implemented in interactive real space refinement, as in Coot (Emsley *et al*., 2010).

**Figure 5.**
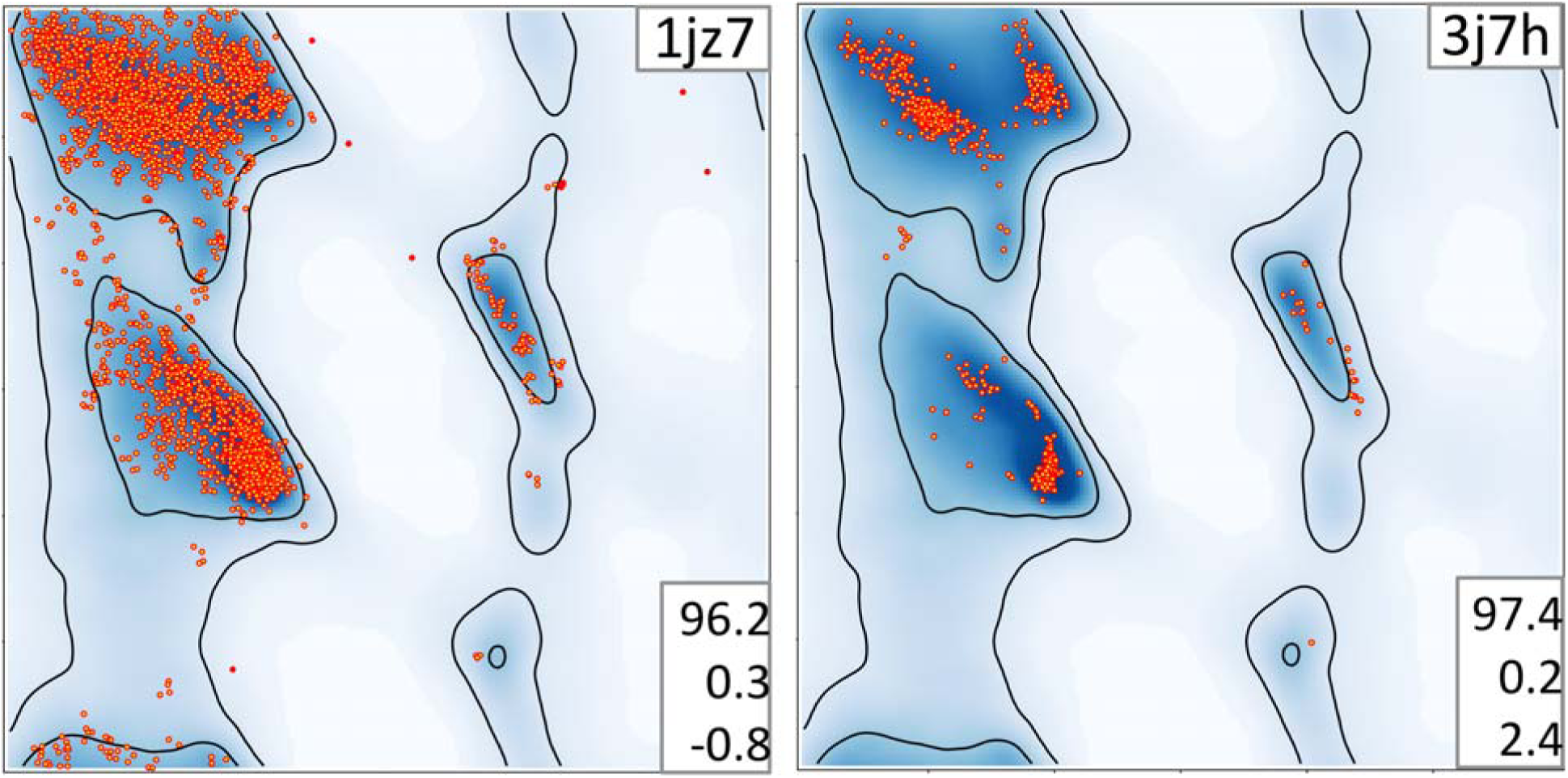
Ramachandran plots for two 100% sequence identical structures: 1jz7 (X-ray, 1.5Å resolution) and 3j7h (cryo-EM, 3.2Å resolution). The cryo-EM structure was refined with Ramachandran plot restraints in Coot (Bartesaghi *et al*., 2014). Triplets of numbers on the bottom right on each plot indicate, from top to bottom: percentage of residues in favored and outlier regions, Rama-Z.

Another example is illustrated in Figure 6. The 3.9Å starting model 5jLh (Fig. 6A) was used to derive 9Å model 6g2t (Fig. 6B), both using cryo-EM data. Comparison of number of residues lies in favored (93.3% and 97.6%) and outlier (1.6% and 0.4%) region would lead us to believe that lower-resolution model has better backbone geometry. However, the high Rama-Z score of 2.5 is in the “suspicious” range compared to the score of −1.6 for 5jLh and suggests that the backbone geometry is worse for 6g2t. Importantly, visual inspection of the 6g2t plot identifies unusual grid-like distributions in the favored regions of the Ramachandran plot. These were not inherited from the starting model (5jLh), so this particular artifact must be the result of the refinement protocol used. That refinement involved H-bond restraints followed by Ramachandran plot restraints (Risi *et al*., 2018). The related models 6cxi and 6cxj have the same pattern in the Ramachandran plot (Suppl. Fig. 3). Apparently, this particular protocol of all-atom refinement systematically produces such artifacts. This leads us to note that a detailed description of the refinement protocol used to obtain final atomic models should always be included in the experimental methods section of structural papers. In these examples, the authors are not “blamed” for producing erroneous Ramachandran features: rather, they are congratulated for describing their experiment in enough detail to help understand the underlying causes and make it possible for developers to create better re-refinement and re-building procedures in the future.

**Figure 6.**
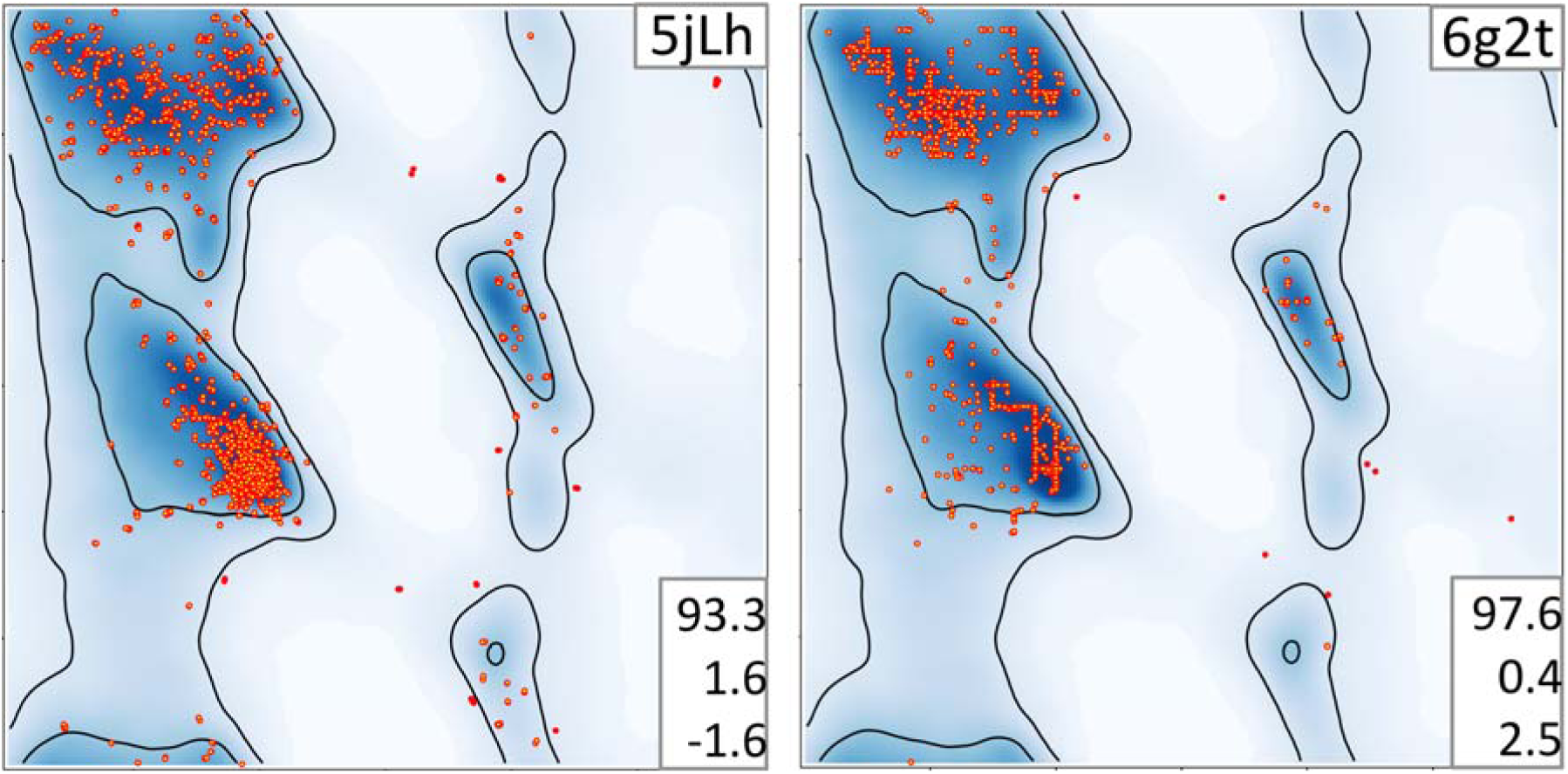
Ramachandran plots of 5jLh used to derive 6g2t. Residues of 6g2t form clear horizontal and vertical lines which is indicated by rather high Rama-Z. Triplets of number on the bottom right on each plot indicate, from top to bottom: percentage of residues in favored and outlier regions, Rama-Z.

### Selected examples from crystallography

In the preceding sections we focused on models obtained using cryo-EM data, as this is a rapidly growing field where there is a predominance of models from lower resolution data. It might be tempting to think that models derived from crystallographic data are less susceptible to the same problems, and that the Rama-Z score has limited applicability. However, analysis of the crystallographic structures in the Protein Data Bank suggests otherwise, as we observe several cases where the score is useful in identifying problematic models. Therefore, the utility and applicability of the Rama-Z validation metric does not depend on the experimental method used to obtain an atomic model. Here we present a number of models with both similar and different artifacts compared to cryo-EM cases to illustrate this point (Fig 7).

**Figure 7.**
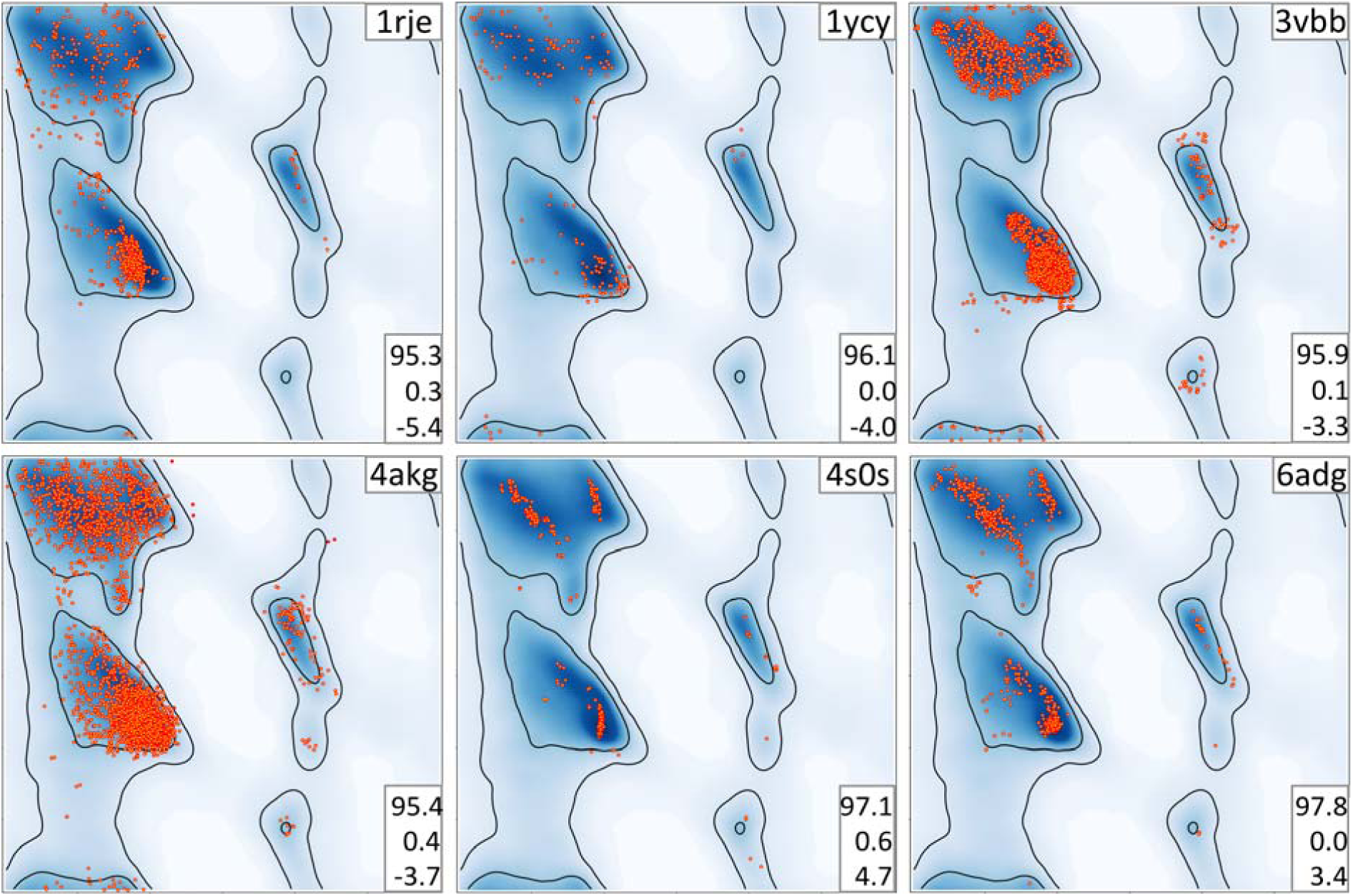
Examples of Ramachandran plots with unusual distribution of (φ, Ψ) angles. Plots are referred to by the PDB code of corresponding atomic model (upper right corner of each plot). Triplets of numbers on the bottom right on each plot indicate, from top to bottom: percentage of residues in favored and outlier regions, Rama-Z.

#### 1rje

This 2Å resolution model (Rfree 0.21) does not show any negative geometry metrics with 95.3% of residues in the favored Ramachandran region and only 0.3% Ramachandran outliers. Nevertheless, an extremely low Rama-Z score of −5.4 indicates something abnormal. Indeed, points on the Ramachandran plot are visibly shifted to the left side of the most favorable alpha-helical region.

#### 1ycy

This lower-resolution (2.8Å) model has 0% Ramachandran outliers. The points in the plot lie around the alpha-helix peak, and only sparsely populate the peak itself. The Rama-Z score of −4 clearly identifies this unusual distribution.

#### 3vbb

Another lower-resolution (2.8Å) model with a low Rama-Z score of −3.3. The Ramachandran distribution displays similar features to structure 6j2q in Figure 4. This reinforces the idea that model artifacts are independent of the type of experiment.

#### 4akg

This 3.3Å model displays too uniform a distribution of residues in the Ramachandran plot without clearly following the most frequent peaks. The abnormality is also indicated by a Rama-Z score of −3.7.

#### 6adg

This model solved with 3Å resolution data shows signs of overfitting with respect to the Ramachandran plot. Most residues occupy the most prominent peaks. The Rama-Z score highlights this abnormality with a very high positive value of 3.4.

#### 4s0s

This model is another example of very high positive Rama-Z score of 4.7. The model was solved with X-ray data at 2.8Å resolution. The Ramachandran plot shows even more overfitting with residues also forming vertical lines in alpha-helical region.

### Limitations of the Rama-Z score

One of the limitations of the Rama-Z score is that it is not very suited for small (sub-) structures with few residues. This is largely a result of the reliance on normalization against a control set of structural models (Hooft *et al*., 1997). In general, normalization is not well suited to small sample sizes, i.e. few available residues. Therefore, the Rama-Z score should be interpreted in light of the calculated uncertainty – the RMSD value; both PDB-REDO and Phenix report these values.

## Conclusions

We have shown that the simple counting of residue fractions that belong to favored and outlier regions of the Ramachandran plot is insufficient to validate protein backbone conformation, particularly when additional restraints have been introduced into the refinement process. These counts may still obey the recommended guidelines, but corresponding Ramachandran plots may show unlikely distributions. These odd distributions may range from trivially identifiable with the naked eye to very subtle. With the an increasing number of lower resolution models becoming available, particularly from cryo-EM, and refinement algorithms actively using all the available information to improve low-resolution refinement by, for example, using Ramachandran plot as restraints, additional validation tools are necessary.

The Ramachandran Z score introduced by Hooft *et al*. more than two decades ago did not make it into mainstream validation procedures, but has now found its place. Here we have demonstrated the utility of this validation metric to pinpoint unlikely distributions of protein main-chain conformations that often are not obvious to an untrained eye nor can be flagged by other standard validation metrics. The expanded database of protein structures has made it possible for us to suggest new cutoffs for Rama-Z validation, with |Rama-Z| > 3 indicating improbable backbone geometry, 2 < |Rama-Z| < 3 unlikely yet possible, and |Rama-Z| < 2 normal backbone geometry. We have also devised a much needed method to calculate the uncertainty in the Rama-Z score, which should be used in its interpretation.

The rapid growth in the number of atomic models derived from lower resolution cryo-EM data, and the concomitant changes in structure refinement algorithms, argues for improved validation metrics. We therefore advocate for greater acceptance of the Rama-Z metric by the structural biology community and note that PDB-REDO has been reporting the Rama-Z score since its inception. Routine use of this metric by researchers refining atomic models at lower resolution, and also by the Protein Data Bank in its validation reports, would likely greatly improve the quality of macromolecular models.

## Availability

The method is implemented and available in open-source CCTBX (*mmtbx*.*rama_z*) library as well as in *Phenix* as a command line tool *phenix*.*rama_z* and also in various validation reports generated by *Phenix*. The *tortoize* implementation is available in PDB-REDO and will become available in the CCP4 and CCP-EM suites in the near future.

## Acknowledgements

This research was supported by the NIH (grant GM063210), the Phenix Industrial Consortium, and by the Netherlands Organization for Scientific Research (NWO; Vidi grant 723.013.003). This work was partially supported by the US Department of Energy under Contract DE-AC02-05CH11231.

## Author contributions

Software: O.V.S., R.P.J., M.L.H.; Investigation: O.V.S., P.V.A., N.W.M., R.P.J., M.L.H.; Writing - Original Draft: O.V.S., P.V.A., N.W.M.; Writing – Review & Editing: A.P., P.D.A.; Supervision: A.P., P.D.A; Funding Acquisition: A.P., R.P.J., P.D.A.

## Footnotes

## Methods

### Rama-Z implementation

The Ramachandran plot Z-score calculation was implemented in the CCTBX closely following the original algorithm from Hooft *et al*. (Hooft *et al*., 1997); a similar re-implementation has been reported for *tortoize* in PDB-REDO (Beusekom, Joosten et. al. 2018). A notable difference is that we used the Top8000 database (Williams *et al*., 2018) of high-quality manually curated models as the underlying data. We used only (φ, Ψ) angles formed by residues with B-factor less than 30Å^2^. A very similar set of models was used to derive the current Ramachandran contours in MolProbity (Williams *et al*., 2018). This led to 1,604,080 residues used in the determination of the distributions. Assignment of secondary structure was performed with the *from_ca* algorithm (Terwilliger *et al*., 2018) with adjusted parameters to make results similar to ones obtained by KSDSSP – an alternative implementation of Kabsch’s & Sander’s DSSP algorithm (Kabsch & Sander, 1983; Joosten, te Beek *et al*., 2011). In addition to the standard residue types we distinguished pre-Proline and *trans*-Proline cases for all secondary structure types and *cis*-Proline for loops. Seleno-methionine residues were counted together with methionine. The total number of residue types was 64: 21 standard residues (including pre-Proline and *trans*-Proline) in the three helix/sheet/loop secondary structures and additional *cis*-Proline in loops. The least populated group is *cis*-Proline with 3953 residues and the most populated group is Glycine in loops with 70,883 residues. The full table of residue counts per group is available in Table S1. The increased number of residues allowed us to reduce the bin size from the original 10° to 4° maintaining a minimum average of one residue per 4° bin for most of the categories. In addition to reporting the whole-model Rama-Z our implementation also reports Rama-Z scores for helices, sheets and strands separately.

Thus, now, in addition to the original implementation in WHAT_CHECK, there are two new implementations: CCTBX and *tortoize* in PDB-REDO.

As there are now multiple implementations that calculate the Rama-Z value, their compatibility should be asserted. The Rama-Z values from CCTBX in Phenix, *tortoize* in PDB-REDO, and WHAT_CHECK were calculated for a test set of 124518 PDB entries. Despite differences between the underlying data and minor technical differences, the correlation between calculated scores, using linear regression, was very high with correlation coefficients of 0.96 between CCTBX and *tortoize*, 0.93 between *CCTBX* and *WHAT_CHECK and 0*.*93 between WHAT_CHECK and tortoize*. Figure S1A shows the relation between CCTBX and tortoize. A slope of linear correlation of 0.8 indicates that the Rama-Z distribution calculated by CCTBX is more “dense” than that of *tortoize*. This is the result of *tortoize* being based on data from the PDB-REDO databank rather than the PDB. We calculated Rama-Z for large number of models and conclude that the latter are very similar, as expected and can be used interchangeably.

### Secondary structure-dependent Rama-Z scores

Separate distributions were calculated using the same method for helices, sheets and loops. If there are enough residues in a respective region, one can get a better insight about the quality of the backbone geometry for secondary structure elements. It should be noted, that *Secondary structure-dependent* Rama-Z scores and the Rama-Z score for the whole model are related in an unobvious way: the scores for helices, sheets, loops and the whole model were calibrated separately to achieve a mean score of 0 and an RMSD of 1 for the reference models, therefore the calibration values are different. This becomes obvious in some corner cases: for example, if the whole model does not have any helices and beta-sheets, the score for loops and for the whole model will be different. As a general guideline we suggest checking the separate the Rama-Z scores when the Rama-Z score for the whole model does not indicate any problems.

### Rama-Z reliability

Since Rama-Z is a statistical metric, the larger the model (more instances of Ramachandran pairs) the smaller the expected error and the more precise the calculated result. To estimate the reliability of the Rama-Z score for a particular model we use the Jackknife method (Quenouille, 1956; Tukey, 1958), resampling to estimate RMSD:

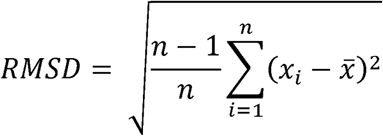

where *n* is the number of (φ, Ψ) pairs in the model,*x*_*i*_ the Rama-Z score calculated for the model with omission of *i*-th (φ, Ψ) pair, 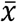 the average of all *x*_*i*_

RMSD values for 143,567 models available in PDB are shown on Figure S1 (B). It can be seen that the RMSD of the Rama-Z score indeed largely depends on the size of the model. Indicatively, for models of 100 residues the average RMSD is 0.73, for models of 1000 residues 0.24, and for models of 5000 residues 0.09. This reliability estimation algorithm has also been implemented in the *tortoize* module of PDB-REDO.

---

**Figure S1.**
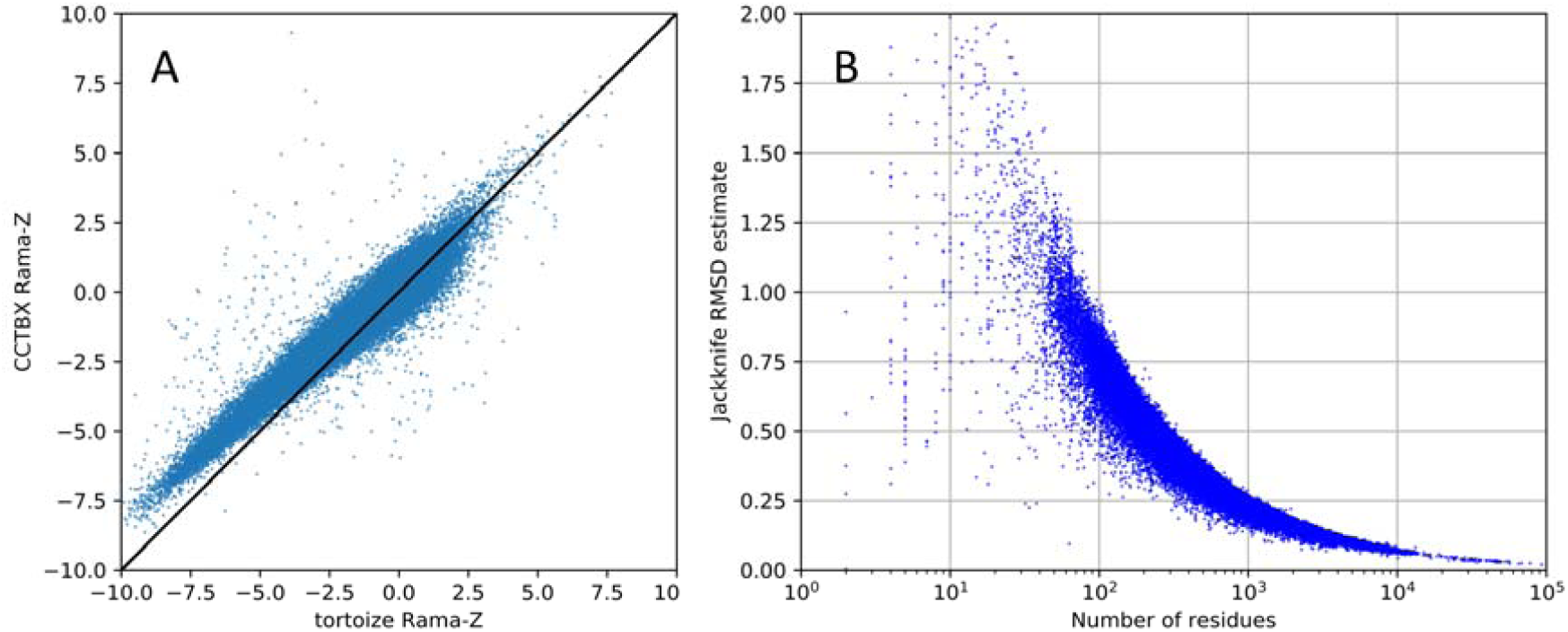
Validating the Rama-Z implementations and RMSD estimation A. Rama-Z from CCTBX vs Rama-Z from *tortoize* for 124518 PDB-REDO entries; the diagonal is marked as a black line. B. Jackknife RMSD estimations (blue dots).

**Figure S2.**
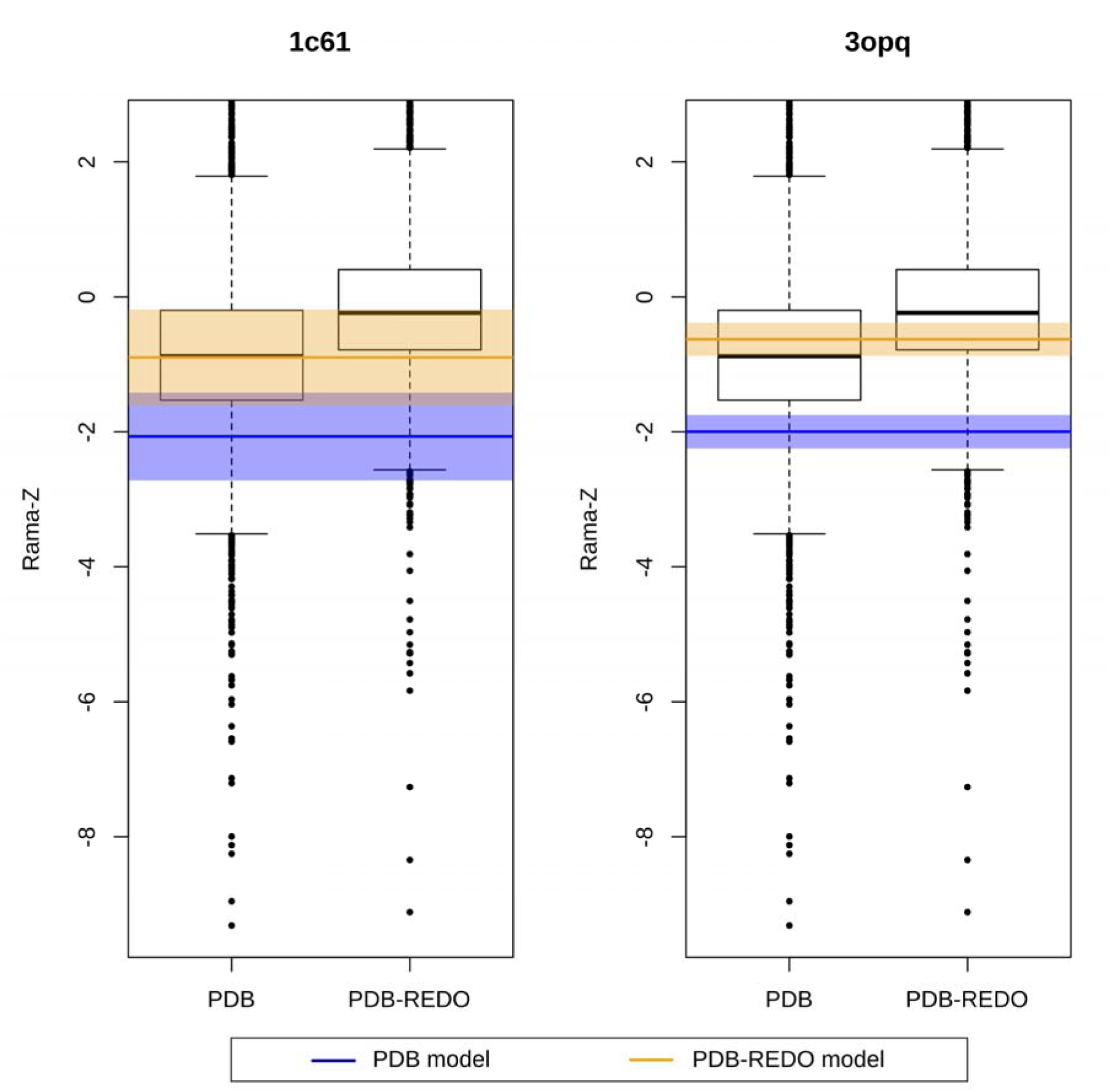
Two examples for the use of the RMSD value of Rama-Z in PDB-REDO: in 1c61 the improvement of the Rama-Z (blue line to orange line) appears significant, but it is not considering the large RMSD for this small structure (blue and orange background, indicating 1= RMSD); in 3opq a similar absolute value change in Rama-Z can be considered significant using the same criterion.

**Figure S3.**
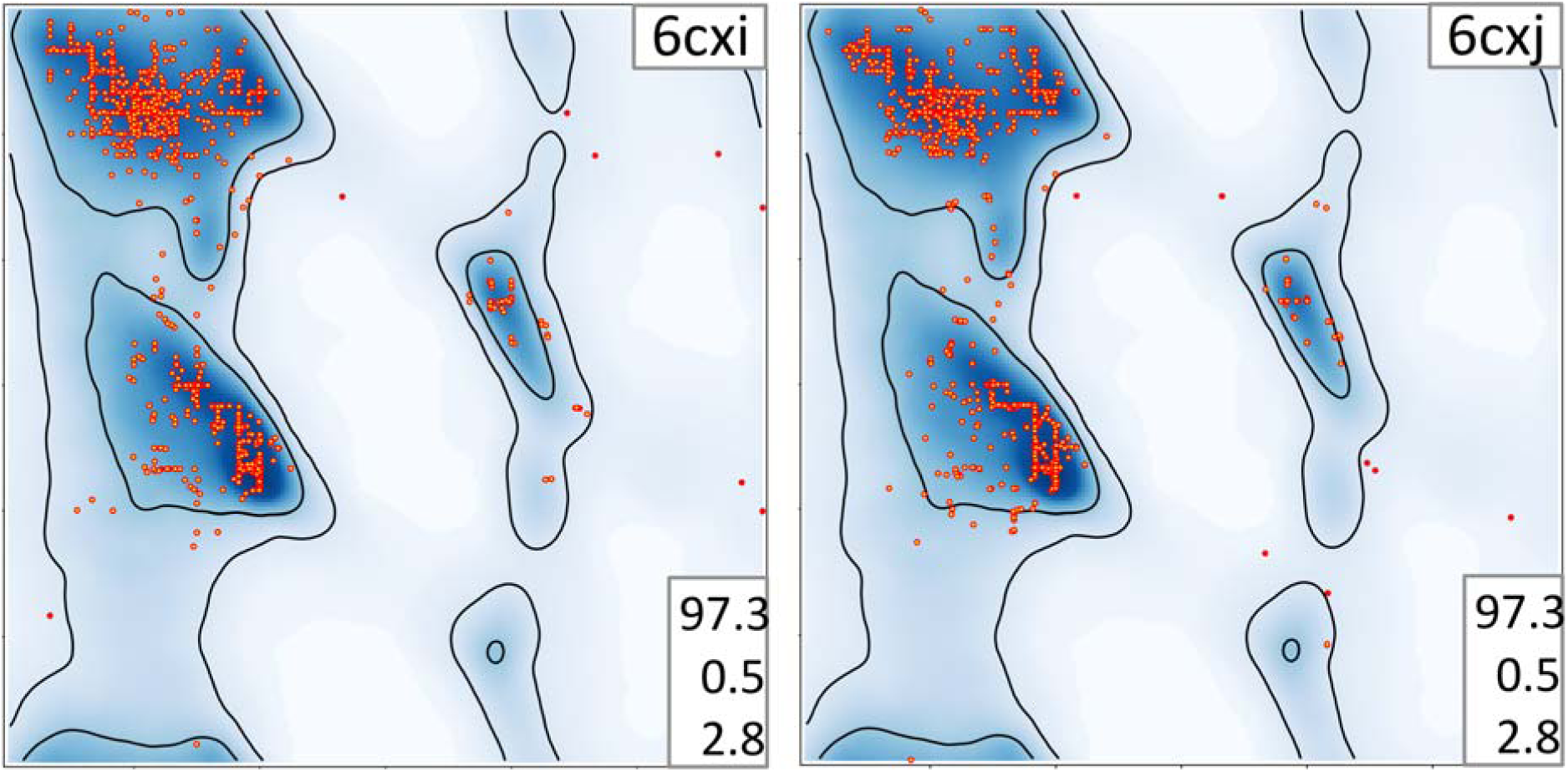
Examples of grid-like distributions on Ramachandran plot with high Rama-Z score. Triplets of numbers on the bottom right on each plot indicate, from top to bottom: percentage of residues in favored and outlier regions, Rama-Z.

**Table S1.** Residue counts.

| Residue type | Helix | Sheet | Loop |
| --- | --- | --- | --- |
| ALA | 70631 | 19896 | 42026 |
| LEU | 67451 | 30469 | 43158 |
| GLU | 49958 | 12828 | 35618 |
| LYS | 36805 | 12588 | 32908 |
| VAL | 35831 | 39369 | 39014 |
| ARG | 35611 | 12931 | 28789 |
| ILE | 33485 | 27816 | 29069 |
| SER | 33288 | 15612 | 41675 |
| ASP | 32091 | 11212 | 43895 |
| GLY | 31498 | 16608 | 70883 |
| THR | 27955 | 19858 | 35816 |
| GLN | 27391 | 7765 | 20117 |
| PHE | 22729 | 16370 | 24794 |
| ASN | 22412 | 8715 | 32360 |
| TYR | 19585 | 13879 | 22883 |
| MET | 16014 | 6738 | 10847 |
| HIS | 12777 | 6802 | 16117 |
| transPRO | 12581 | 7615 | 47244 |
| prePRO | 12093 | 8164 | 44056 |
| TRP | 8341 | 5154 | 9754 |
| CYS | 6606 | 5211 | 8371 |
| cisPRO | - | - | 3953 |

## Footnotes

1 The usage of “Ramachandran plot” refers to the list of (φ, Ψ) values from a specific model being superposed on the contoured heatmap representation of the expected (φ, Ψ) values determined from a filtered set of accurate proteins.

2 PDB protein codes follow the convention outlined in Moriarty (2015)

